# The deubiquitylase USP9X controls ribosomal stalling

**DOI:** 10.1101/2020.04.15.042291

**Authors:** Anne Clancy, Claire Heride, Adán Pinto-Fernández, Andreas Kallinos, Katherine J. Kayser-Bricker, Weiping Wang, Victoria Smith, Hannah Elcocks, Simon Davis, Shawn Fessler, Crystal McKinnon, Marie Katz, Tim Hammonds, Neil P. Jones, Jonathan O’Connell, Bruce Follows, Steven Mischke, Justin A. Caravella, Stephanos Ioannidis, Christopher Dinsmore, Sunkyu Kim, Axel Behrens, David Komander, Benedikt M. Kessler, Sylvie Urbé, Michael J. Clague

## Abstract

When a ribosome stalls during translation, it runs the risk of collision with a trailing ribosome. Such an encounter leads to the formation of a stable di-ribosome complex, which needs to be resolved by a dedicated machinery. The initial stalling and the subsequent resolution of di-ribosomal complexes requires activity of Makorin and ZNF598 ubiquitin E3 ligases respectively, through ubiquitylation of the eS10 and uS10 sub-units of the ribosome. It is common for the stability of RING E3 ligases to be regulated by an interacting deubiquitylase (DUB), which often opposes auto-ubiquitylation of the E3. Here, we show that the DUB USP9X directly interacts with ZNF598 and regulates its abundance through the control of protein stability in human cells. We have developed a highly specific small molecule inhibitor of USP9X. Proteomics analysis, following inhibitor treatment of HCT116 cells, confirms previous reports linking USP9X with centrosome associated protein stability and reveals loss of ZNF598 and Makorin 2. In the absence of USP9X or following chemical inhibition of its catalytic activity, steady state levels of Makorins and ZNF598 are diminished and the ribosomal quality control pathway is impaired.

## Introduction

Prompt sensing and resolution of aberrant protein translation is essential for the maintenance of protein homeostasis. Several circumstances can give rise to stalled ribosomes, such as insufficiency of a cognate acylated-tRNA, defective mRNA or faulty ribosomes [1, 2]. The most common cause of ribosomal stalling is thought to be the translation of poly(A), when a nascent mRNA is inappropriately polyadenylated within its coding region to generate a “non-stop” mRNA transcript lacking a stop codon [3, 4]. If a ribosome stalls during translation it risks being rear-ended by a trailing ribosome. This collision generates a stable di-ribosome complex with a defined structure, which is resolved by the engagement of a dedicated machinery [5, 6]. In such cases the E3-ligase ZNF598 ubiquitylates 40S complexes at specific sites on eS10 and uS10 sub-units at the di-ribosome interface [5, 7–9]. This prevents further translation and initiates quality control processes (e.g. degradation of the associated mRNA) and ribosomal recycling pathways through partially understood mechanisms [10]. ZNF598 is a human RING domain protein that shares homology with the yeast protein Hel2, the deletion of which promotes read-through of polybasic sequences [8, 11]. A recent report has provided evidence that the E3-ligases Makorin 1 (MKRN1) and potentially Makorin 2 (MKRN2) may complement the activity of ZNF598 in the ribosomal quality control pathway, by promoting the initial stalling of the leading ribosome as it encounters a polyA tract [12].

The function of E3-ligases, can be opposed by ~100 deubiquitylase (DUB) enzymes drawn from seven families [13]. RING E3s show a propensity to auto-ubiquitylate, leading to their destabilisation, which can be rescued by the activity of interacting DUBs. The best known such example is provided by the association between USP7 and MDM2, which has made USP7 a prominent drug target, as a means to regulate levels of p53 [14]. Recent work, focused on this enzyme, has established proof of principle that selective small molecule inhibition amongst the USP family can be achieved [15–18]. USP9X is one of the most abundant members of the USP family and has been linked with many processes, including centrosome function, chromosome alignment during mitosis, EGF receptor degradation, chemo-sensitisation and circadian rhythms [19–23]. Loss of function mutations in females lead to congenital malformations and intellectual disability [24].

In this study we identify the E3 ligase ZNF598 as a USP9X binding partner and show that USP9X governs the stability of ZNF598. The loss or inhibition of USP9X leads to a substantive reduction in steady state levels of ZNF598 and Makorins that disables an effective response to the presence of stalled ribosomes.

## Results and discussion

In a large scale proteomic study of the ribosome interactome, USP9X was the only DUB family member to be identified [25]. Furthermore, USP9X is also apparent within the set of ZNF598 interacting proteins, previously identified in a label free proteomic study [7]. We sought to confirm this interaction by immuno-precipitating FLAG-tagged ZNF598 transiently expressed in HEK293T cells (Figure 1). USP9X is clearly present in the IP containing ZNF598-FLAG and is absent from control lanes.

**Figure 1:**
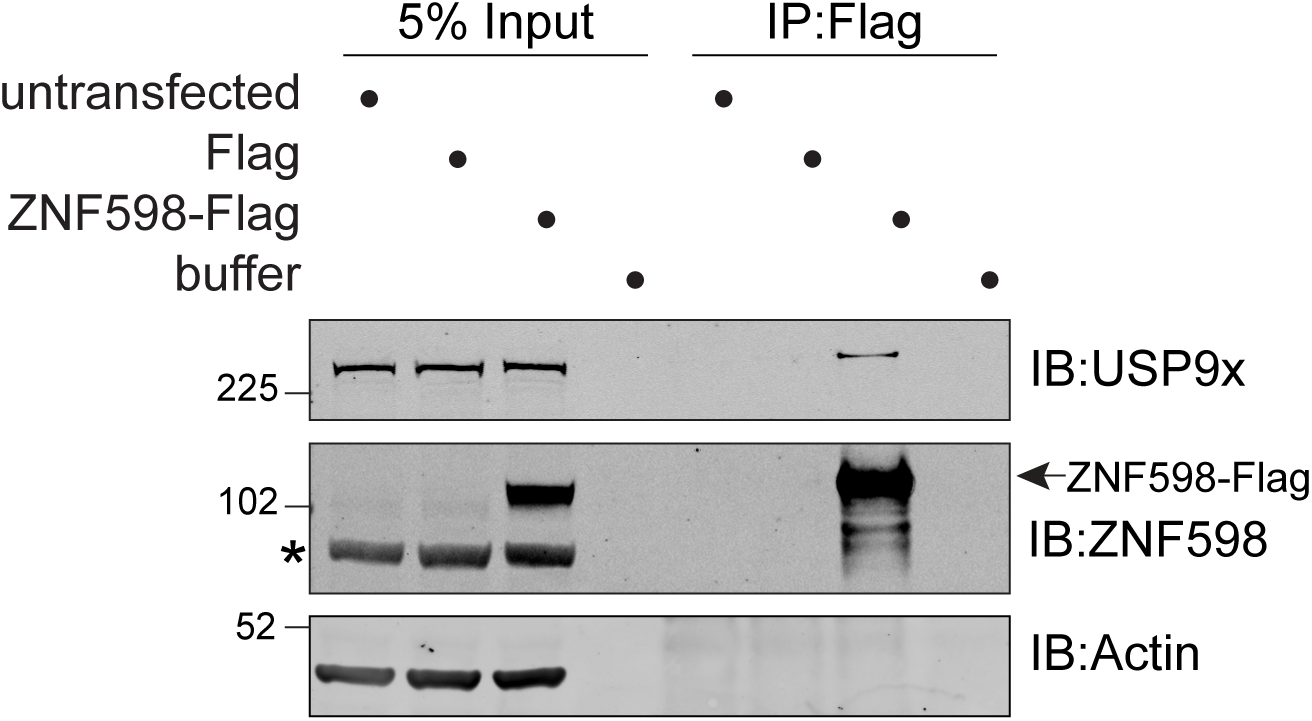
USP9X co-immunoprecipitates with Flag-tagged ZNF598. HEK293T cells were transfected with ZNF598-Flag or Flag alone (pCMV-Tag2B) and cell lysates were subjected to immunoprecipitation (IP) with Flag-antibody coupled agarose beads. IPs were probed alongside 5% of the input as indicated. Arrowhead indicates ZNF598-Flag; asterisk indicates a non-specific band.

We next compared the engineered USP9X^-/0^ HCT116 colon cancer cells that have been described previously [19] with wild type cells of the same origin. As expected, the USP9X^-/0^ cells show reduced levels of previously identified peri-centrosomal substrates CEP55, CEP131 and PCM1 [21, 22] (Figure 2A). ZNF598 levels are also greatly diminished in these cells (Figure 2A and Supplementary Figure 1). Two lines of argument suggest that this is not an effect on transcription, (i) endogenous ZNF598 mRNA levels are similar between the two cell lines (Figure 2B) and (ii) levels of exogenous HA-ZNF598 expression that is driven by a non-native promoter are also diminished in transfected cells (Figure 2C). We next treated these cells with cycloheximide and monitored the decay of the expressed HA-ZNF598. In wild type cells, levels of HA-ZNF598 remained stable over the 6 hours of incubation, whilst in the USP9X^-/0^ cells the levels significantly decline to about ~60% in the same period (Figure 2D). The most parsimonious explanation of these combined results is that USP9X interacts with ZNF598 and regulates its steady state levels through the control of its stability.

**Figure 2:**
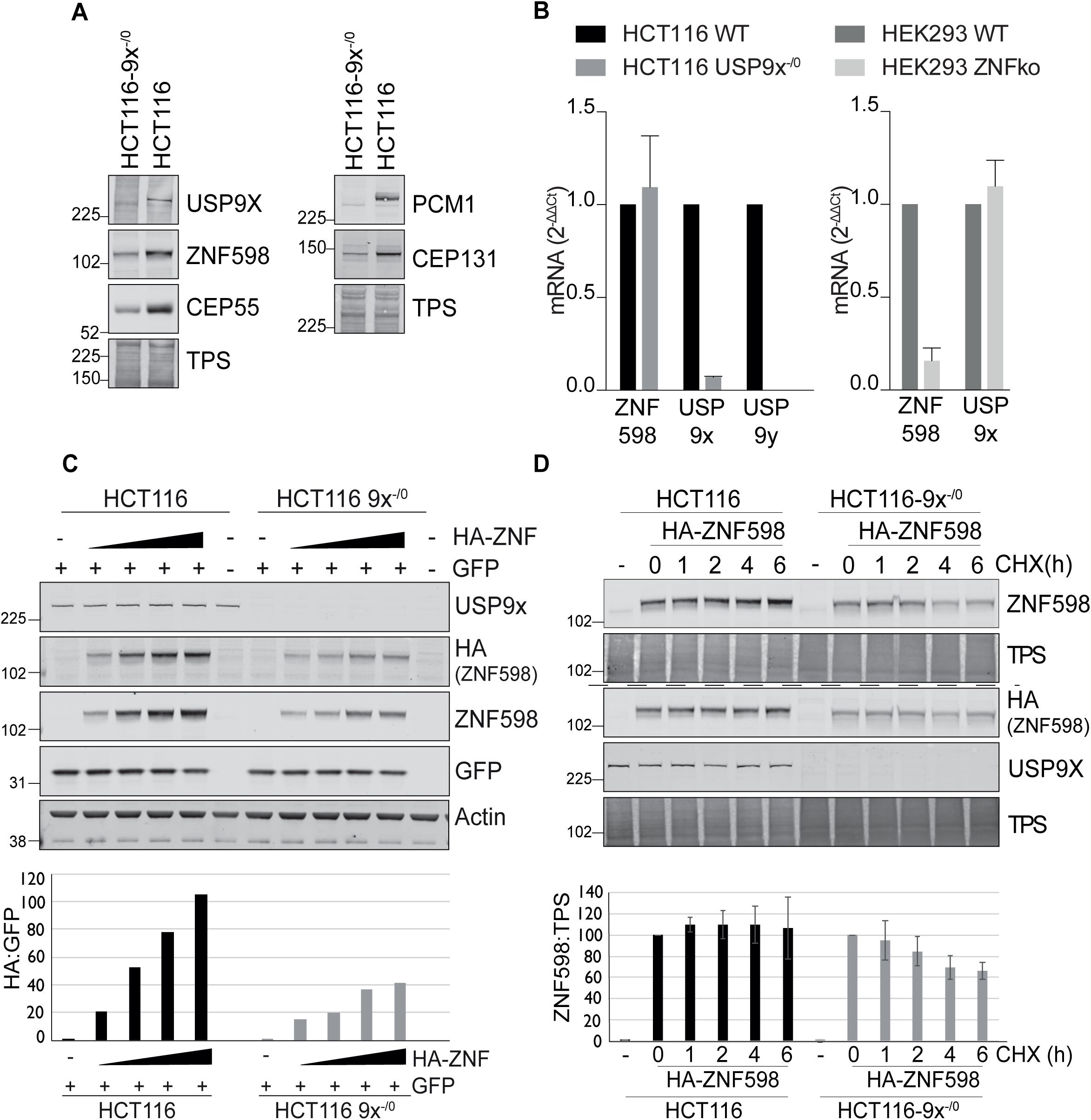
ZNF598 is destabilised in USP9X KO cells. **A -** HCT116 or HCT116 USP9x^-/0^ lysates were analysed by immunoblot with the indicated antibodies. **B -** qRT-PCR reactions for ZNF598, USP9X and USP9Y (normalised to Actin) were performed with cDNA derived from the indicated cell lines. The mean of three independent biological replicates is shown and error bars indicate the standard deviation. **C -** HCT116 or HCT116 USP9x^-/0^ cells were transfected with 0, 0.2, 0.4, 0.8 or 1.6μg HA-ZNF598 and 0.2μg GFP as a transfection control, and lysates analysed by immunoblot with the indicated antibodies. Graph shows HA-ZNF598 relative to co-transfected GFP. **D -** HCT116 or HCT116 USP9x^−/0^ cells were transfected with 0.2μg HA ZNF598 and treated for the indicated times with 100μg/ml Cycloheximide (CHX). Lysates (8µg for HCT116 and 20µg HCT116-USP9x^−/0^) were probed with the indicated antibodies. Graph shows the average of 4 independent experiments, error bars represent the standard deviation. TPS: Total Protein Stain.

To demonstrate that this requires the catalytic activity of USP9X we took advantage of a highly selective small molecule inhibitor FT709 (Figure 3A, chemical structure, 3B-3H). We identified USP9X inhibitors using a ubiquitin-TAMRA fluorescence polarization HTS assay, screening the inhibitory potential of a diverse collection of approximately 140,000 compounds available at FORMA Therapeutics. Primary hits were further validated for direct USP9X binding by biophysical techniques such as surface plasmon resonance (SPR). Optimization of hits with respect to activity and physicochemical properties resulted in a series of compounds that included FT709.

**Figure 3:**
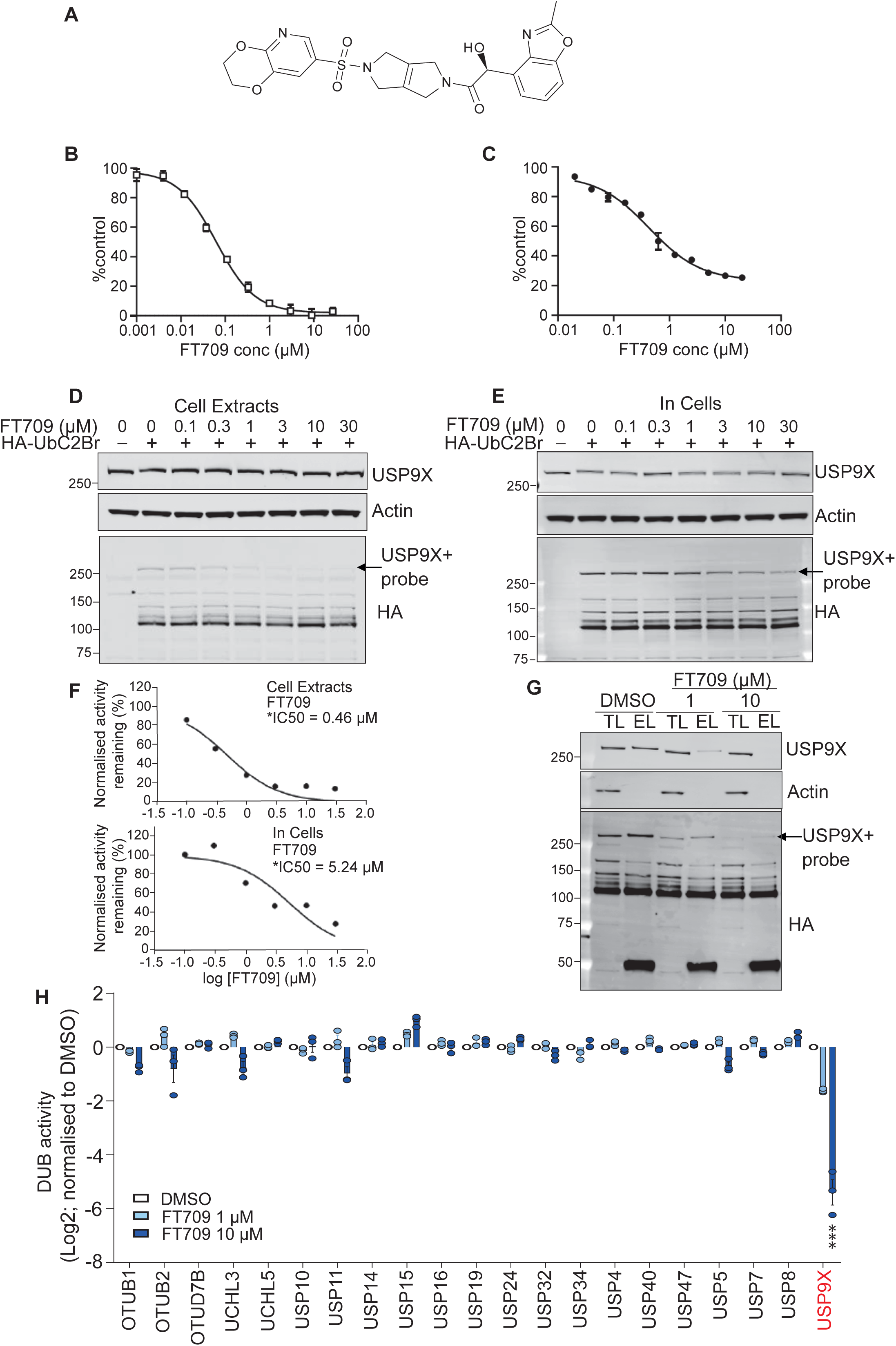
Characterisation of a highly selective USP9X inhibitor. **A -** Chemical structure of FT709. **B** *In vitro* potency of FT709 against USP9X. **C** BXPC3 cell-based potency of FT709 for reduction of CEP55. **D, E, F -** Cell lysates (D) or intact MCF7 cells (E) were incubated with FT709 (30 minutes at 25°C for cell extracts, 3 hours at 37°C for cells) at the indicated concentrations. Cells were lysed, and extracts incubated with 0.1 µg HA–UbC2Br probe for 5 min at 37°C, followed by SDS-PAGE analysis. Samples were immunoblotted with USP9X and HA antibodies as indicated. Arrow indicates HA-probe labelled band corresponding to the USP9X~Ub probe adduct. Modification of USP9X with a ubiquitin probe (USP9X~Ub) was lost with increasing concentrations of inhibitor. **F -** quantitation of Western blots **G, H -** HA-based immunoprecipitation of HA-UbC2Br probe labelled DUBs from cell lysates incubated first with DMSO, 1 or 10 µM FT709 for one hour at 37°C. Immuno-precipitated proteins were eluted and either analysed side by side with total lysate samples by immunoblotting (TL; Total lysate; EL: Eluate) or were subjected to mass spectrometry-based quantification in three technical replicates. Differences in DUB-probe binding were quantified for 23 identified DUBs and normalised relative to DMSO control. FT709 only affects USP9X (Dunnett multiple comparisons test; ***: p<0.001; see Methods for statistics and Supplementary Figure 2 for uncropped immunoblots).

FT709 is potent in a biochemical assay with an IC_50_ of 82 nM (Fig 3B). Modulation of CEP55 expression in BxPC3 pancreatic cancer cells showed an IC_50_ of 131 nM (Fig 3C). The selectivity of FT709 was tested across more than 20 DUBs in a biochemical assay (Table 1) and was inactive across the panel (IC50>25 µM). FT709 shows vastly improved specificity over the compound WP1130, which has been previously used as a USP9X inhibitor tool compound [26, 27]. FT709 competes with an active site probe (HA-UbC2Br) with an IC_50_ of ~0.5µM when applied to MCF7 breast cancer cell lysates and ~5µM to intact MCF7 cells (Figure 3D-F and Supplementary Figure 2). Immunoprecipitation from cell lysates labelled with the active site probe HA-UbC2Br, revealed that USP9X is uniquely sensitive to this compound, within a set of 23 DUBs quantified by mass spectrometry (Figure 3G,H and Supplementary Figure 2).

**Table 1.**

| DUB | IC50<br>(μM) |
| --- | --- |
| USP1 | >25.8 |
| USP2 | >26.7 |
| USP4 | >33.3 |
| USP5 | >33.3 |
| USP7 | >25.8 |
| USP8 | >25 |
| USP9x | 0.082 |
| USP10 | >33.3 |
| USP11 | >33.3 |
| USP12 | >33.3 |
| USP13 | >26.7 |
| USP15 | >26.7 |
| USP18 | >27.9 |
| USP22 | >26.7 |
| USP24 | >26.7 |
| USP25 | >26.7 |
| USP28 | >26.7 |
| USP33 | >26.7 |
| USP36 | >26.7 |
| USP37 | >33.3 |
| USP47 | >33.3 |

Acute inhibition with FT709, recapitulates gene deletion of USP9X in HCT116 cells, leading to reduction of ZNF598 (Figure 4A). We conducted a wider survey of protein expression following USP9X inhibition through quantitative mass spectrometry (Figure 4B, Supplementary Table 1). Amongst a small number of proteins that decrease by more than two-fold following inhibitor treatment, known USP9X substrates are prominent. These include the (peri)-centrosomal proteins PCM1, CEP55 and CEP131 [21, 22] and the mitotic kinase, TTK, also known as Monopolar spindle 1 (Mps1) kinase (Figure 4B,C) [28]. ZNF598 is also found within this cohort, in alignment with our Western blot analysis (Figure 4A,B). Intriguingly a second RING E3-ligase MKRN2, that has been linked to ribosome stalling, is is equally identified as a clear outlier (Figure 4B, C) [12]. Accordingly MKRN2 is also reduced in USP9X^-/0^ cells (Figure 4D). Note that these are the only RING E3 ligases, contained within the proteomic dataset (>6000 proteins), which show this magnitude of response to USP9X inhibition (Supplementary Table 1).

**Figure 4:**
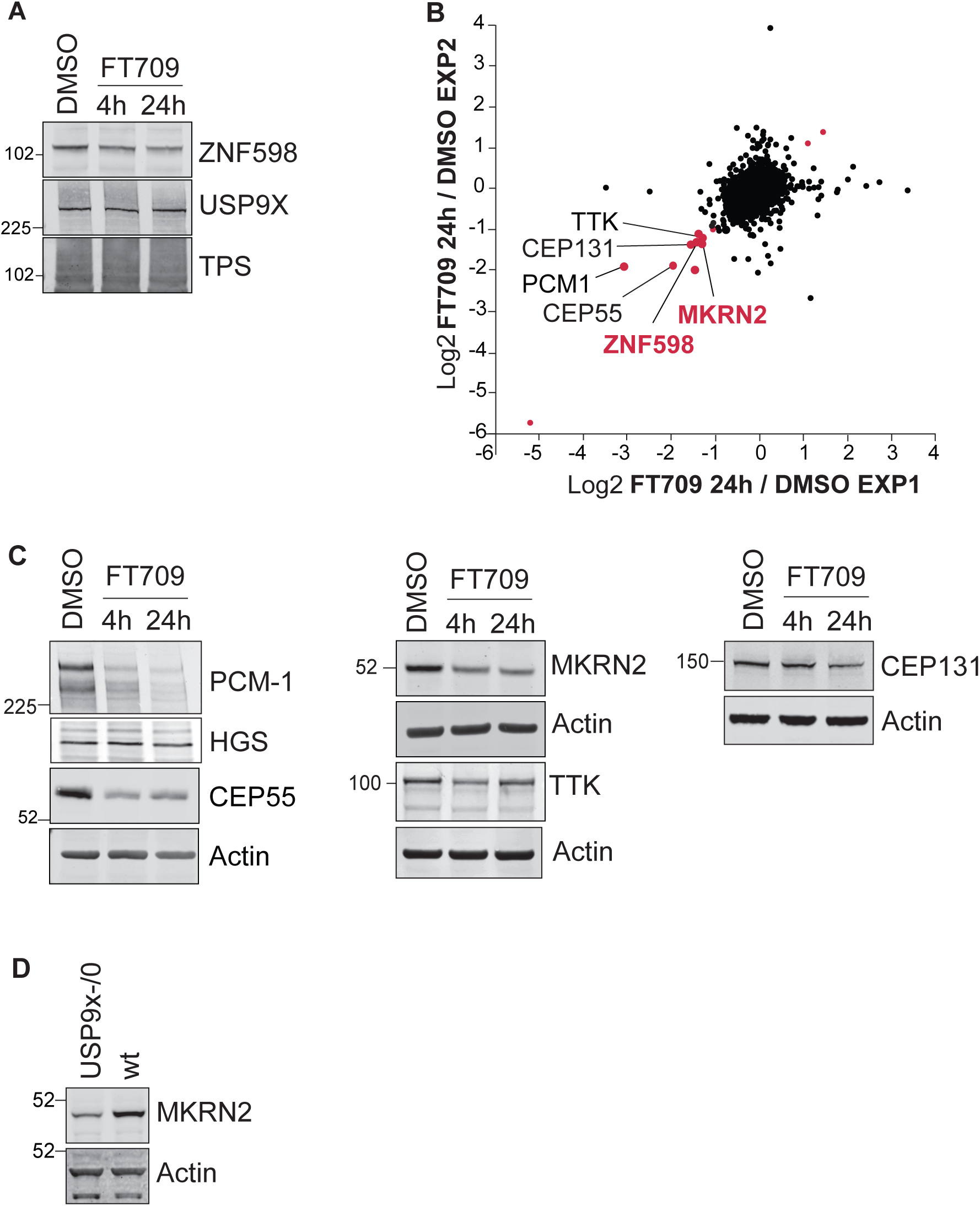
Inhibition of USP9X catalytic activity depletes ZNF598 and MKRN2 protein levels. **A -** HCT116 cells were treated for 4 or 24 hours with a selective USP9X inhibitor (FT709, 10µM). Cells were lysed in RIPA buffer and samples analysed by SDS-PAGE and immunoblotted for ZNF598 and USP9X. **B -** Correlation of two distinct experimental SILAC based proteomic datasets showing the de-enrichment of ZNF598 and MKRN2 alongside known USP9X substrates (in black type) in HCT116 cells treated for 24 h with USP9X inhibitor (FT709, 10 µM). Outliers for which the ratio is either lower than Log_2_(−1.0) or larger than Log_2_(+1) in both datasets are shown in red. **C -** Western blot validation of USP9X inhibitor sensitive proteins identified in B. HCT116 were treated with FT709 at 5 µM (CEP131), or 10 µM (all other samples) and analysed as in A. TPS: Total Protein Stain. **D** -De-enrichment of MKRN2 in USP9X knockout cells. HCT116 or HCT116 USP9x^-/0^ lysates were analysed by immunoblot with the indicated antibodies.

We next asked if the effects of USP9X ablation upon ZNF598 and MKRN2 could be extended to another cell type, HEK293 cells. We used a set of 4 individual gRNAs designed to target USP9X, which were packaged in an expression plasmid that also codes for Cas9 (px459-pSpCas9(BB)-2A-Puro-v2). Plasmids were transfected either individually, or as a pool, and cell populations were harvested after 168 hours of selection with puromycin. The pooled transfection effectively ablated USP9X protein expression across the cell population, leading to a correspondingly stark reduction in ZNF598 and MKRN2 levels. ZNF598 loss, achieved through the same transient CRISPR/Cas9 based approach or in a stable knock-out cell line showed no reciprocal effect on USP9X levels (Figure 5A).

The HEK293 cells used in this study (HEK293-Flp-IN TREX GFP-P2A-(K^AAA^)_21_ -P2A-RFP) have been engineered to express a reporter system for terminal ribosomal stalling [8]. The reporter cassette contains GFP (N-terminal) and RFP (C-terminal) separated by a FLAG-tagged stalling reporter (SR) incorporating a polyA stretch of twenty-one codons (K^AAA^)_21_ (Figure 5B). This is flanked by viral P2A sequences, at which ribosomes skip formation of a peptide bond, without interrupting translation elongation. Consequently, unimpaired translation generates 3 proteins (GFP, FLAG-SR, RFP) in equal amounts. Stalling at the FLAG-SR aborts translation prior to RFP synthesis, leading to a sub-stoichiometric RFP:GFP ratio. Failure to effectively respond to stalled ribosomes allows eventual read-through and a consequent rise in the RFP:GFP ratio that can be assessed by fluorescence activated cell sorting (Figure 5B, schematic diagram)[8]. As reported previously, an isogenic reporter cell line, in which the ZNF598 gene has been deleted, shows an enhanced RFP:GFP ratio when compared to parental cells consistent with read-through, due to failure of the ribosomal stalling response [8]. Using the pooled USP9X gRNA cells, that show highly reduced levels of both USP9X and ZNF598, we could recapitulate this phenomenon. We also analysed 3 of the cell populations treated with individual gRNAs as represented in Fig 5A that show varying effectiveness for USP9X depletion. Guide 1 serves as a control because it was ineffective in editing the USP9X gene and accordingly shows no change in the RFP:GFP ratio. Guide 4 generates two distinct populations with about 50% showing an enhanced RFP:GFP ratio, whilst Guide 7 shows a uniform enhancement of this ratio reminiscent of the effect of ZNF598 deletion and comparable to the pooled gRNA sample (Figure 5C). Importantly, we were able to recapitulate polyA read through with FT709, implicating USP9X enzymatic activity, and providing a pharmaceutical approach to counteract ribosomal stalling (Figure 5D). In this condition, effects on ZNF598 levels were less dramatic in this cell line, but were accompanied by losses of both MKRN1 and MKRN2. We propose that USP9X inhibition may promote read through by a combined effect on each of these 3 RING E3 ligases linked to the ribosomal quality control pathway (Figure 5E).

**Figure 5:**
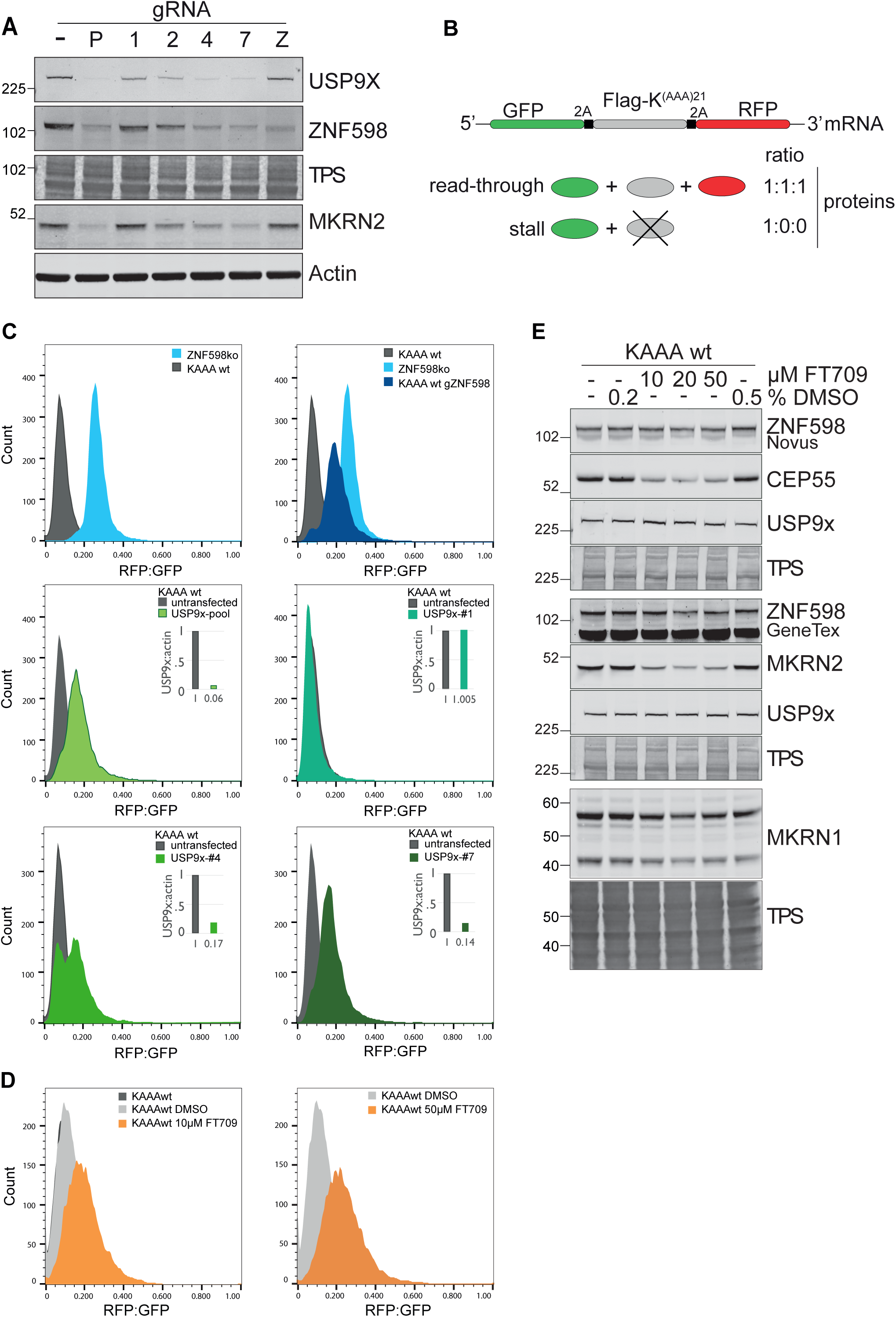
USP9X ablation or inhibition impairs the ribosomal stalling response. **A -** HEK293 Flp-In T-Rex GFP-P2A-(KAAA)21-P2A-RFP wt cells (KAAA wt) were transfected with a plasmid containing Cas9 and gRNAs targeting USP9X or ZNF598. P = pool of USP9X guides, 1/2/4/7= individual USP9X guides, Z = ZNF598 guide. Lysates were analysed 168 hrs after transfection and selection in Puromycin, by immunoblotting with the indicated antibodies. **B -** Schematic of the fluorescent ribosomal stalling reporter expressed in this cell line. If stalling is not efficiently resolved, read-through occurs, the Flag-SR and RFP are expressed. **C -** FACS analysis of the RFP:GFP ratio in KAAA wt or ZNF598ko cells following transfection with px459-pSpCas9(BB)-2A-Puro_v2 containing gRNA as indicated. Cells were gated for live singlets, then for GFP positive cells. Insets indicate the USP9X protein levels normalised to wt untransfected cells. **D -** FACS analysis of the RFP:GFP ratio in wt cells following inhibition of USP9X by FT709 for 72 hrs. **E** - KAAA wt cells were treated with indicated concentrations of FT709 as in D and analysed by immunoblotting with selected antibodies. TPS: Total Protein Stain.

## Conclusions

USP9X is for the most part a non-essential DUB family member, that is nevertheless expressed at relatively high levels [29, 30]. Multiple biological functions have been ascribed to USP9X that include roles in apoptosis, Wnt signaling and mitotic check-point control [31, 32] [33]. Our proteomics data most strongly support previously established links to centrosome biology [21, 22, 34]. Here, we reveal a new biological role for USP9X in the resolution of stalled ribosomes, which is supported by unbiased proteomics. We propose that this is principally related to its governance of ZNF598 stability, but could also extend to Makorin family members [12]. It is possible that USP9X could also play a more direct role in the deubiquitylation of ribosomal sub-units themselves during stalling resolution. However this is difficult to unravel from effects upon their ubiquitin conjugation by the E3 ligases described here. Moreover, other DUBs (USP21, OTUD3) have recently been linked with this function [35]. The non-uniform dynamics of ribosomal processing, duration and resolution of stalling may have important implications for protein folding, mRNA turnover and for the integrated stress response (ISR) [6, 36]. Recent studies have also shown that ribosomal collisions can result in +1 frame-shifting when the no-go RNA decay pathway is compromised [37]. Our introduction of a highly specific USP9X tool compound inhibitor will enable further enquiry into pathways previously linked to USP9X, which should now include global profiling of protein translation.

## Methods

### Chemical compound

FT709 used in this study was prepared by the procedures described in detail previously [38]. Cell culture HEK293T cells were cultured in DMEM with GlutaMAX+ 10% tetracycline-free FBS, HEK293 Flp-IN 293 Trex K^(AAA)^21 wt or ZNF598ko were cultured in DMEM with GlutaMAX+ 10% tetracycline-free FBS 15 μg/ml blasticidin, 100 μg/ml hygromycin. To induce reporter expression 1µgml^-1^ Doxycycline was added 24 hr before harvesting. HCT116 and HCT116 USP9x^-/0^ were cultured in McCoys media + 10% FBS. MCF7 cells were cultured in DMEM medium supplemented with 10% FCS, 1% penicillin/streptomycin and 1% glutamine at 37°C and 5% CO2. Cells were routinely checked for mycoplasma. For cycloheximide assay, cells were treated for the indicated times with 100µg/ml cycloheximide and harvested 24 hr post transfection.

### Transfection

For transient transfection, 2µg total DNA (per well of a 6 well plate) was transfected using Genejuice (Novagen) according to manufacturer’s instructions. Cells were harvested 24-168 hr post transfection.

### Lysis and Western Blot analysis

Cultured cells were lysed for 10 min at 4°C in RIPA buffer (10 mM Tris–HCl pH 7.5, 150 mM NaCl, 1% Triton X-100, 0.1% SDS, 1% sodium deoxycholate) supplemented with mammalian protease inhibitor cocktail (SIGMA). Proteins were resolved using SDS–PAGE (Invitrogen NuPage gel 4–12%), transferred to nitrocellulose membrane, blocked in 5% fat-free milk or 5% bovine serum albumin in TBS supplemented with Tween-20, and probed with primary antibodies overnight. Visualisation and quantification of Western blots were performed using IRdye 800CW and 680LT coupled secondary antibodies and an Odyssey infrared scanner (LI-COR Biosciences, Lincoln, NE).

### Co-IP

Cells were lysed in TNTE buffer (10mM Tris-Cl pH 7.5, 150mM NaCl 0.3% Triton-X100, 5mM EDTA) supplemented with mammalian protease inhibitor cocktail (Sigma). 750µg total protein was then incubated with 20µg prewashed FLAG affinity gel (Sigma) for 2h at 4°C, washed with TBS buffer (10mM Tris-Cl pH 7.5, 100mM NaCl) and eluted in sample buffer (62.5mM Tris-Cl pH 6.8, 3% SDS, 10% glycerol, 3.2% ß-mercaptoethanol). Immunoprecipitates were then analysed by western blot as above.

### RNA isolation and Reverse Transcription qPCR

Total RNA was isolated from HCT116, HCT116 USP9x^−/0^, Flp-IN 293 Trex K(^AAA^)21 wt and Flp-IN 293 Trex K(^AAA^)21 ZNF598 KO using the Qiagen RNA extraction kit (74106). cDNA was generated using 1 µg RNA and the Thermo Scientific RevertAir H Minus reverse transcriptase (Fisher Scientific UK: 11541515) supplemented with RNasin (Promega:N251S), PCR nucleotide mix (Promega: U144B) and oligo (dT) 15 Primer (Promega: C1101). qPCRs were performed in triplicate using primers against ZNF598, USP9x, USP9y, actin (see Table 2 for sequences) and iTaq Mastermix (BioRad: 172-5171) in a BioRad CFX Connect real time system. The mean Ct values were normalised to actin (ΔCt=Ct target - Ct actin), raised to the exponent of 2^−ΔCt^ and normalised to the respective wild-type control cell line to generate 2^−ΔΔCt^.

**Table 2.**
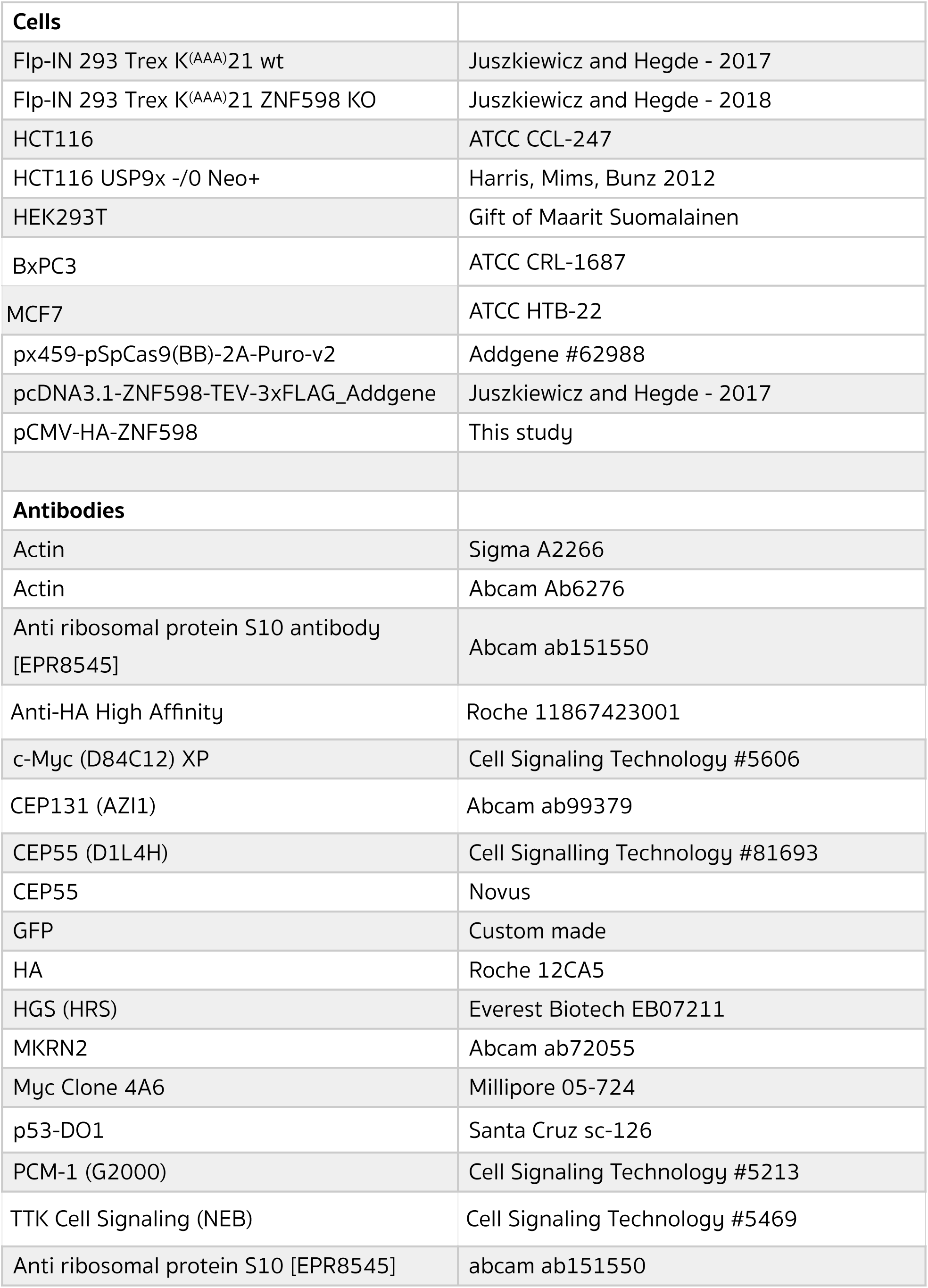
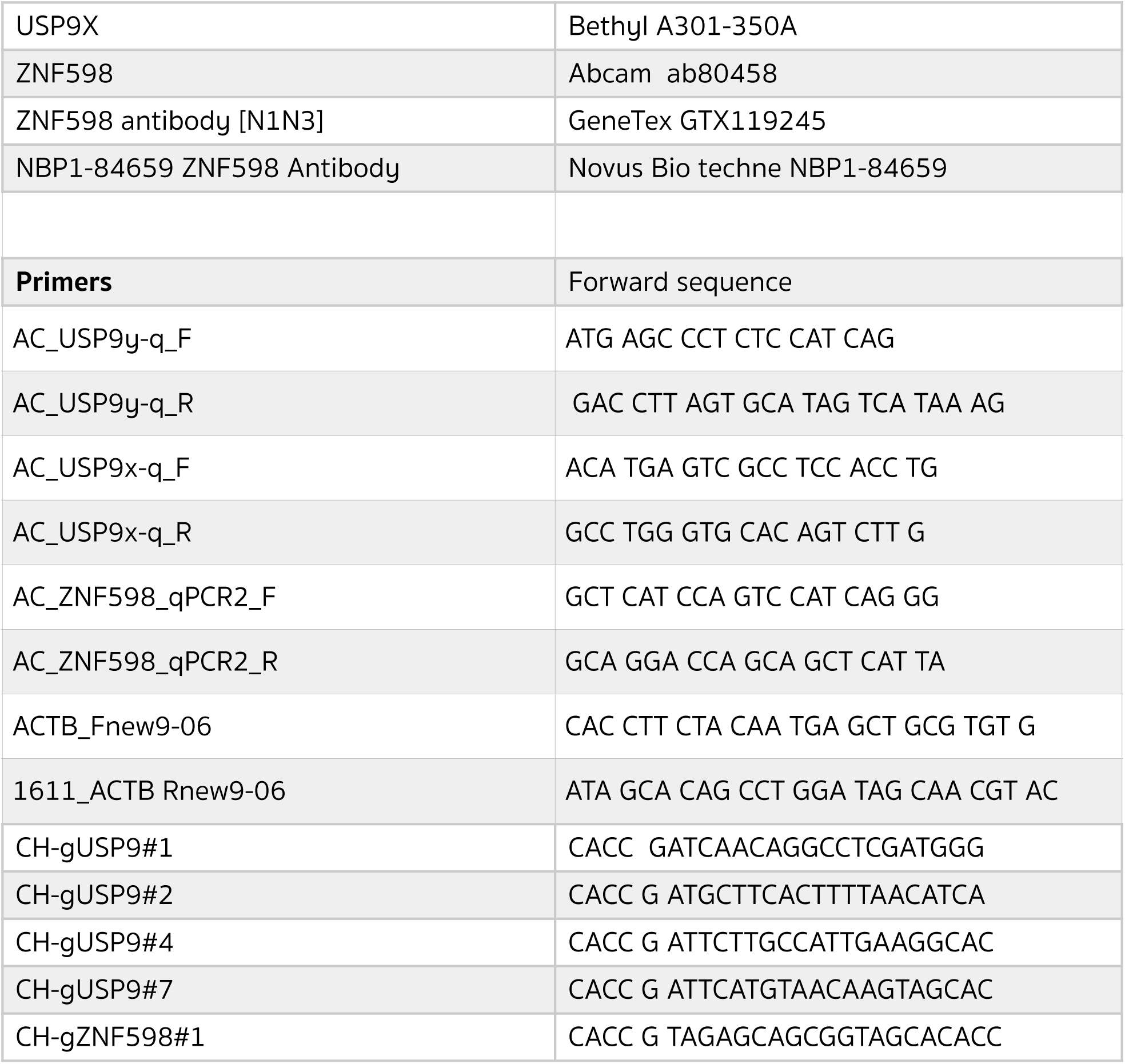
List of reagents used in this study

### FACS

Primers as detailed in Table 2 were cloned into px459-pSpCas9(BB)-2A-Puro-v2 vector at the BbsI site. These were transfected into Flp-IN 293 Trex K(AAA)21 wt and ZNF598 ko using Genejuice (Novagen). 24 hours post-transfection, media was changed and 1µg/ml Puromycin included. Cells were then cultured for 7 days before harvesting for western blot and FACS analysis. For FACS, cells were trypsinised, counted, resuspended in 10% tetracycline free FBS in PBS and analysed on a FACS Aria III in conjunction with FlowJo software.

### DUB biochemical assay

The assay was performed in a final volume of 6 µL in assay buffer containing 20 mM Tris buffer pH8, 0.03% bovine γ globulin, 0.01% Triton X-100 and 1 mM glutathione. Nanoliter quantities of 10-point, 3-fold serial dilutions in DMSO were pre-dispensed into 1536 assay plates for a final top concentration between 25 to 33.3mM and subsequent half-log dilutions. 2X DUB (0.025nM final concentration) was added and pre-incubated for 30 minutes at room temperature. 2X Ub-Rhodamine (25 nM final concentration) was added to initiate the reaction. Fluorescence readings (Ex: 485 nm, Em: 535 nm) were acquired over 12 minutes (Envision reader). The slope of the data from each point was used to determine IC_50_.

For all assays, data were reported as percent inhibition compared with control wells. IC_50_ values were determined by curve fitting of the standard 4 parameter logistic fitting algorithm included in the Activity Base software package: IDBS XE Designer Model205. Data were fitted using the Levenburg Marquardt algorithm. Results presented are based on 3 independent experiments performed in quadruplicates.

### MSD assay for CEP55

BxPC-3 cells were seeded in 96-well plates and exposed to FT709 (20 µM top concentration, 1:2 serial dilutions) for 6 hours. Cell lysates were prepared in RIPA buffer and stored at −80°C until analysis. Samples were analysed by an MSD Elisa assay (Pacific Biolabs) using a CEP55 antibody (Novux, 1:500 dilution in PBS) captured overnight at 4°C, 30µL lysates per well, 30 µL of CEP55 Ab (CST#81693) diluted 1:2000 in 1% blocker A/PBS and 30 uL per well of a 1:4000 diluted Goat anti-Rabbit Sulfo-tag, 1% Blocker A/PBS. Plates were read on a MSD Sector Imager 2400. Results were transformed as % DMSO controls and curves fitted using a non-linear regression to determine the IC50. Results presented are based on duplicate values.

### DUB profiling assays using Ub-based active site directed probes

Molecular probes based on the ubiquitin scaffold were generated and used essentially as described [39, 40]. In brief, HA-tagged Ubiquitin bromoethyl (HA-UbC2Br) was synthesised by expressing the fusion protein HA-Ub_75_-Intein-Chitin binding domain in E.Coli BL21 strains [41]. Bacterial lysates were prepared and the fusion protein purified over a chitin binding column (NEB labs, UK). HA-Ub_75_-thioester was obtained by incubating the column material with mercaptosulfonate sodium salt (MESNa) overnight at 37°C. HA-Ub_75_-thioester was concentrated to a concentration of ~1mg/ml using 3,000 MW filters (Sartorius) and then desalted against PBS using a PD10 column (GE Healthcare). 500 μL of 1-2mg/mL of HA-Ub_75_-thiolester was incubated with 0.2mmol of bromo-ethylamine at pH 8-9 for 20 minutes at room temperature, followed by a desalting step against phosphate buffer pH 8 as described above. Ub probe material was concentrated to ~1mg/ml, using 3,000 MW filters (Sartorius), and kept as aliquots at −80°C until use.

### DUB competition assays with cell extracts and with cells (in situ)

Crude MCF7 cell extracts were prepared as described previously using glass-bead lysis in 50mM Tris pH 7.4, 5mM MgCl_2_, 0.5mM EDTA, 250mM sucrose, 1mM DTT [40, 41]. For experiments with crude cell extracts, 50μg of MCF7 cell lysate was incubated with different concentrations of FT709 for one hour at 37°C, followed by addition of 1μg HA-UbC2Br and incubation for 5 minutes at 37°C. Incubation with Ub-probe was optimised to minimise replacement of non-covalent inhibitor FT709 by the covalent probe. Samples were then subsequently boiled in reducing SDS-sample buffer, separated by SDS-PAGE and analysed by Western Blotting using anti-HA (1:2000), anti-USP9x (CST, 1:1000) or beta Actin (1:2000) antibodies. 5×10^6^ intact MCF7 cells were incubated with different concentrations of inhibitors in cultured medium for 4 hours at 37°C, followed by glass-bead lysis, labelling with HA-UbC2Br probe and analysis by SDS-PAGE and Western blotting as described above.

### Mass spectrometry based DUB inhibitor profiling assays

Ub-probe pulldown experiments in presence of different concentrations of the inhibitors FT709 were performed essentially as described [39, 40] with some modifications. In brief, immuno-precipitated material from 500μg^-1^mg of MCF-7 cell crude extract was subjected to in-solution trypsin digestion and desalted using C18 SepPak cartidges (Waters) based on the manufacturer’s instructions. Digested samples were analyzed by nano-UPLC-MS/MS using a Dionex Ultimate 3000 nano UPLC with EASY spray column (75μm x 500 mm, 2μm particle size, Thermo Scientific) with a 60 minute gradient of 0.1% formic acid in 5% DMSO to 0.1% formic acid to 35% acetonitrile in 5% DMSO at a flow rate of ~250nl/min (~600bar/40°C column temperature). MS data was acquired with an Orbitrap Q Exactive High Field (HF) instrument in which survey scans were acquired at a resolution of 60.000 @ 400m/z and the 20 most abundant precursors were selected for CID fragmentation. From raw MS files, peak list files were generated with MSConvert (Proteowizard V3.0.5211) using the 200 most abundant peaks/spectrum. The Mascot (V2.3, Matrix Science) search engine was used for protein identification at a false discovery rate of 1%, mass deviation of 10ppm for MS1 0.06 Da (Q Exactive HF) for MS2 spectra, cys carbamidylation as fixed modification, met oxidation and Gln deamidation as variable modification. Searches were performed against the UniProtKB human sequence data base (retrieved 15.10.2014). Label-free quantitation was performed using MaxQuant Software (version 1.5.3.8), and data further analysed using GraphPad Prism software (v7) and Microsoft Excel. Statistical test-s ANOVA (multiple comparison; Original FRD method of Benjamini and Hochberg) was performed using GraphPad Prism software [42].

### SILAC based proteome analysis of FT709-treated HCT116 cells

HCT116 cells were grown in SILAC DMEM supplemented with 10 % dialysed FBS (Dundee Cell Products) at 37°C and 5 % CO2. To generate light, medium and heavy stable isotope-labelled cells, arginine- and lysine-free DMEM medium was supplemented with 200 mg/L L-proline and either L-lysine (Lys0) together with L-arginine (Arg0) (“Light”), L-lysine-2H_4_ (Lys4) with L-arginine-U-13C_6_ (Arg6) (“Medium”) or L-lysine-U-13C_6_-15N_2_ (Lys8) with L-arginine-U-13C_6_-15N_4_ (Arg10) (“Heavy”) at final concentrations of 84 mg/L for the arginine and 146 mg/L for the lysine until fully metabolically labelled. Cells were treated with DMSO or 10 µM FT709 for 4 hours or 24 hours, prior to lysis in 50 mM Tris pH 6.8, 2 % SDS, 10 % glycerol. Relative protein concentrations of the lysates were determined using a BCA-assay (Thermo-Fisher) and “Light”, “Medium” and “Heavy” labelled lysates were combined in a 1:1:1 ratio.

### Deep proteome workflow

Protein extracts (1.2 to 1.5 mg) containing SDS were reduced with 5 mM dithiothreitol, alkylated with 20 mM iodoacetamide and then subjected to methanol/chloroform extraction. Protein pellets were resuspended in 6 M urea by vortexing and sonication, then diluted to a final concentration of 1 M prior to in-solution digestion with 0.2 µg/µl trypsin (Sequencing grade (Promega) overnight at 37°C. Off-line high-pH reverse-phase prefractionation was performed on the digested material as previously described [43], with the exception that eluted peptides were concatenated down to 10 fractions. Peptide fractions were analysed in technical replicates by nano-UPLC-MS/MS using a Dionex Ultimate 3000 nano UPLC with EASY spray column (75 μm x 500 mm, 2 μm particle size, Thermo Scientific) with a 60 minute gradient of 2% acetonitrile, 0.1% formic acid in 5% DMSO to 35% acetonitrile, 0.1% formic acid in 5% DMSO at a flow rate of ~250nl/min. MS data was acquired with an Orbitrap Q Exactive HF instrument in which survey scans were acquired at a resolution of 60.000 at 200m/z and the 20 most abundant precursors were selected for HCD fragmentation with a normalised collision energy of 28.

### Data analysis

All raw MS files from the biological replicates of the SILAC-proteome experiments were processed with the MaxQuant software suite; version 1.5.3.8 using the Uniprot database (uniprotHumanUP000005640.fasta-retrieved in July 2015) and the default settings [44]. The minimum required peptide length was set to 6 amino acids and two missed cleavages were allowed. Cysteine carbamidomethylation was set as a fixed modification, whereas oxidation and acetyl N terminal were considered as variable modifications. ProteinGroup text files were further processed using Excel (see Supplementary Table 1) and the log_2_ of the normalised ratios were plotted using JMP software (version 13.0.0).

## Acknowledgements

We thank Ramanujan Hegde (LMB, Cambridge) for his provision of ribosomal stalling reporter assay reagents and Fred Bunz (John Hopkins University) for sharing USP9X knock-out cells.

---

**Supplementary Figure 1.**
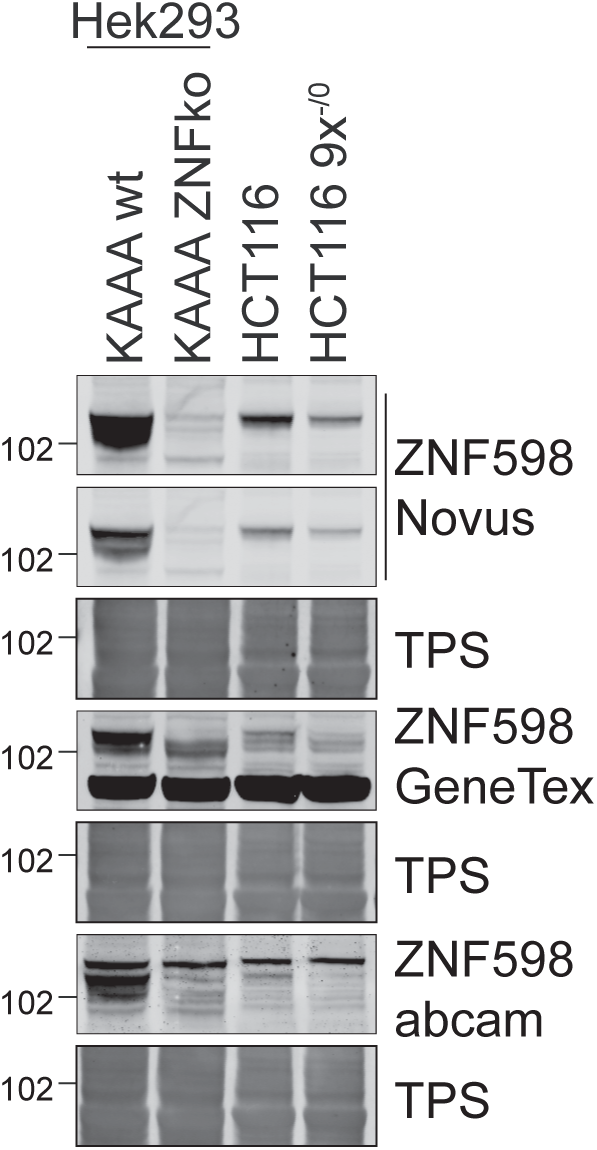
ZNF598 antibody validation. HEK293 Flp-In T-Rex GFP-P2A-(KAA)21-P2A-RFP wt or ZNF598 knockout (ko), and HCT116 wt or HCT116 USP9x^-/0^ cell lysates were analysed by immunoblotting with the indicated antibodies.

**Supplementary Figure 2.**
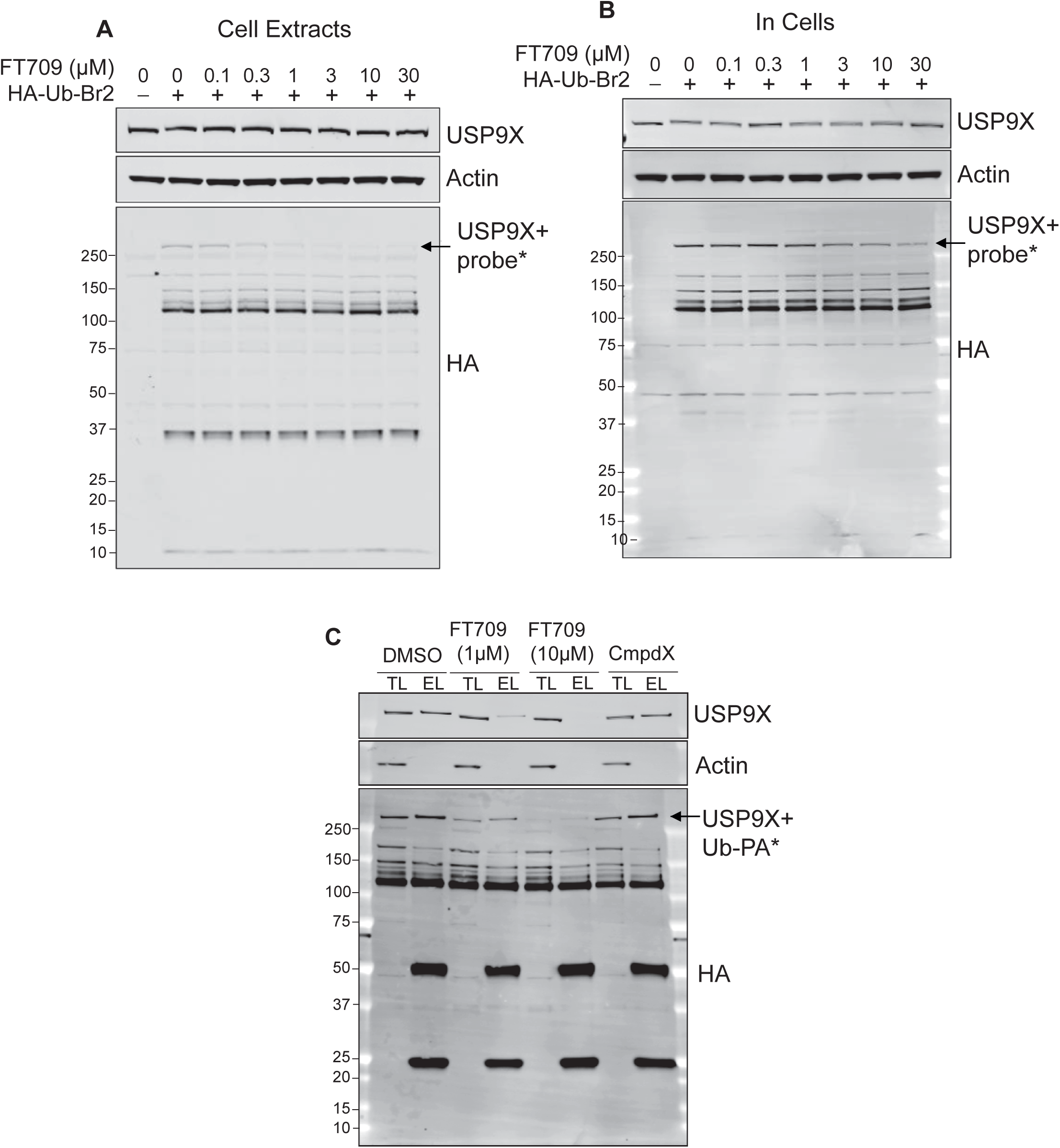
Full western blots of panels D, E and G shown in Figure 3.

